# From Decellurization to Imaging to 3-D Printing: low-cost plant-derived 3D-printed tissue scaffolds for tissue engineering

**DOI:** 10.1101/2020.12.31.424959

**Authors:** Mariam Imran, Sarah Khan, Faisal F. Khan

## Abstract

Nature’s most abundant carbohydrate, cellulose, has incredible structural properties that can be leveraged as scaffolds for tissue engineering. With plants being an inexpensive and easily accessible source, it is more feasible to experiment with these techniques and progress in the field of regenerative tissue engineering. In this study, we set out to optimize a low-cost method to obtain cellulose scaffolds that could potentially mimic a blood vessels after recellularization with endothelial cells. We chose a readily available plant specimen, i.e. cauliflower stalk, which offers anatomical similarity to blood vessels, vascular architecture and interconnected porosity. We went on to capture the cellulose scaffold digitally and created a 3-D model using a computer-aided design (CAD) software which was then used for 3-D printing the scaffold in two different sizes. We believe the decellularize-image-print cycle allows for skipping decellularization processes of new scaffolds every time a scaffold is required and therefore cutting cost and time needed, enables instant dissemination between individual researchers and communities and allows scalable printing at any size and level of detail. We hope this will catalyze even faster innovation in the space of tissue engineering and regenerative medicine.

## I. INTRODUCTION

The demand for organs for transplantation is far greater than their availability. This makes organ transplantation a non-sustainable option to cater for the global demand and patients are in a dire need of an alternative. In the US alone, more than 100,000 patients are on the donor waiting list at any given time and an average of 22 people die every day waiting [1]. One promising alternative that attempts to help in the bioengineering of tissues and organs is the use of recellularized biocompatible scaffolds. Plants then become a subject of interest as they offer a substantial source of a variety of scaffolds. There is an existing use-case for cellulose in a variety of applications in regenerative medicine already including bones [2,3], cartilage [4] and wound healing [5] which is another reason that makes plants an interesting source. Apart from the durability and architectural hierarchy [6] of these scaffolds, they are also not prone to digestion by mammalian cells due to the absence of cellulase, which are all added benefits.

Current research in this domain focuses on the decellularization of plants and recellularization of the remaining scaffold with mammalian cells, with the ultimate aim of transplantation into the human body. Scientists at the Pelling Lab have created prototypes of human ears from apples [7,8]. *Ficus Hispida, Pachira aquatica* and a species of *Garcinia* plant types were made fit for recellularization with mammalian cells using two protocols mentioned by Adamski and team [9]. In another experiment, spinach leaves were converted into scaffolds that were later recellularized by human cells [10]. It is also possible to remove cellular material from a donor’s organ and recellularize it with the patient’s cells. Once the cellular material is removed, it results in a non-immunogenic decellularized graft [11]. This can later be recellularized to create an autologous graft [12]. However, mammalian organs are not only short in supply to begin with, but are also expensive. Cellulose scaffolds are a much cheaper alternative.

The advent of 3D printing technology in 1980s revolutionized the additive manufacturing industry. A solid object is formed through deposition of successive layers of an acrylic-based photopolymer which is simultaneously crosslinked by UV light. After adoption of this tool by the biomedical community, bioengineers began experimenting with ‘bio-inks’ which were mainly polymers of biological origin which had the right properties for potential use as an ink in a bioprinter, instead of a plastic. The new 3D ‘bio’ printing variant used bioink comprising of biomaterials deposited layer by layer to form a tissue or an organ [19].

In this research, our goal is to find an optimal low-cost method of decellularization that removes maximum cellular material from a plant tissue and 2) to demonstrate how computer-aided design (CAD) and 3D printing methodologies can be leveraged to innovate faster in the tissue engineering space. We applied multiple protocols and eventually developed a hybrid protocol that was suitable for the specimen taken in this study. We also imaged and captured the details of the decellularized plant-derived scaffold and converted it into a design that was 3D-printer-friendly. The scaffold was then printed in two different sizes to demonstrate how such a library of CAD models are not only readily shareable for dissemination in the community but also scalable and printable in different sizes as per the need. All this, without the need to go back to nature, identify a specimen and optimize and repeat the entire decellularization effort.

3-D Modeling of the obtained cellulose scaffold is one of the most fascinating and promising aspects of this project. Not only can we “steal” innumerable types of scaffolds from nature, we can image them and create 3-D models to share with others. This skips the entire cost and time required for the decellularization process by any subsequent experiment. We can envisage the creation of a variety of libraries online for and by scientists and DIY biologists alike, to choose the most appropriate scaffold designs for their 3-D bio-printing experiments, speeding up the rate of innovation in the field. The scalability aspect of the obtained has an added bonus; in theory, we could create a capillary blood vessel for a human or an elephant from the same scaffold.

The manuscript is organized as follows: Section II presents different methods of decellularization, Section III describes the processes we followed. A proposed hybrid protocol is presented in Section IV, followed by a histological analysis of the treated specimens in Section V. The results of the histological analysis are discussed in Section VI. Confirmed decellularized scaffolds are imaged, modelled and 3-D printed as mentioned in Section VII.

## II. MATERIALS AND METHODS

The plant specimen we selected in this study was a stalk from the head or curd of a white cauliflower (*Brassica oleracea*), due to its high-level anatomical resemblance to a blood vessel, vascular architecture, interconnected porosity and accessibility in the market. The internal region of the cauliflower stalk was scraped out to resemble a human capillary. We tested four different protocols for the decellularization processes from existing literature [7-10]. Table 1 compares the four protocols in terms of methods, materials, any deviations we made and their final result. Based on all four iterations, we propose and develop a hybrid protocol that was the most suitable for the plant specimen used in our study.

**Table 1:** Protocols of Decellularization

| Protocols | A | B | C | D |
| --- | --- | --- | --- | --- |
| Ref | Modeulvsky 2015 [8] | Adamski 2018 (A) [9] | Adamski 2018 (B) [9] | Gershlak 2017 [10] |
| Materials | • 1% SDS Solution | • 5% Bleach solution<br>• 3% NaHCO<sub>3</sub> | • 10% SDS solution<br>• 5% Bleach solution<br>• 1% Triton X-100 | • Hexanes<br>• 1 x PBS |
| Methods | • Wash with distilled water<br>• Submerge in 1% SDS solution | • Wash<br>• Put on low shake<br>• Warm Bleach & NaHCO<sub>3</sub> solution to 70°C<br>• Put in solution at low shake | • Wash<br>• Place in 10% SDS solution<br>• Submerge in 1% Triton X-100 and 5% bleach solution on low shake | • Place cauliflower pieces in n-hexane and 1 x PBS solution<br>• Following steps are same as Protocol C |
| Deviation | None | None | Duration of 10% SDS soak was 10 days instead of 5<br>5% Bleach solution instead of 10% | Steps following n-hexane and 1 x PBS soak abandoned |
| End Color | Light Green | White | White | Light Green |

## III. CONDUCT OF PROTOCOLS

In this section, we describe each of the four protocols, listed in Table 1 in detail. We discuss each of them in detail and present the experimental results.

### Protocol A

Following the decellularization protocol outlined in [8], cauliflower pieces measuring 3cm x 1cm were washed using distilled water to remove debris. They were then submerged in distilled water for 5 minutes. As mentioned in the protocol, 1% SDS solution can be used for better results instead of dish soap. Later, 1% SDS solution was prepared and cauliflower pieces were submerged in it at room temperature. The solution was replaced with a fresh 1% SDS solution every 4-6 hours to prevent it from becoming unsterile due to bacterial growth. After three days, the cauliflower stalks looked significantly lighter and were preserved in isopropanol [13]. The process is illustrated in Fig. 1, with each day’s progress and preservation at the end.

**Fig 1:**
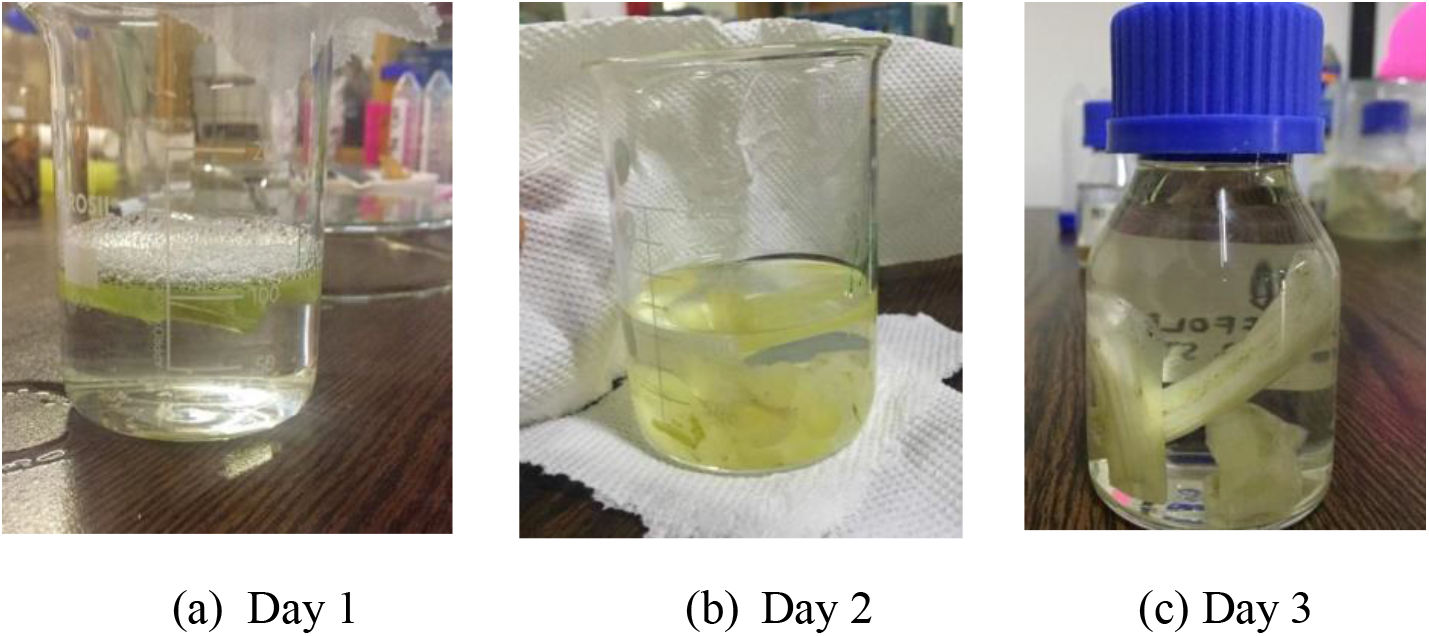
Cauliflower stalk in 1% SDS solution each day and preserved

### Protocol B

The second decellularization technique is a detergent-free protocol [9]. We used a solution of Bleach and Sodium Bicarbonate. Earlier work performed as early as 17th century demonstrated that this solution aids in the separation of the vasculature from the surrounding soft tissue, contributing to the decellularization process [14]. In this protocol, two cauliflower pieces, 2.7 cm x 1.2 cm and 0.8 cm x 0.5 cm, were washed using distilled water to remove debris and were subsequently submerged in distilled water at room temperature. It was then put on low sake on a DIY shake plate, as shown in Fig. 2, made from a vortex, a large petri plate and tissue paper. A solution was prepared using equal volumes of 5% Bleach (NaClO) and 3% Sodium Bicarbonate. The solution was then warmed to 70°C using a water bath. Cauliflower pieces were submerged in the solution and put on low shake. In less than an hour the pieces began to turn white as shown in Fig. 3.

**Fig 2:**
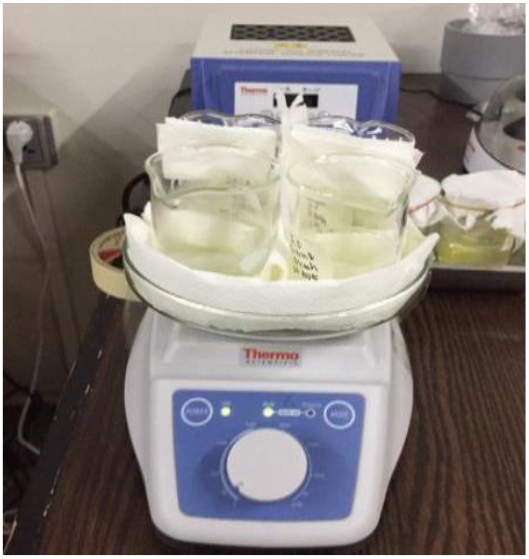
DIY Shake Plate with samples loaded.

**Fig 3:**
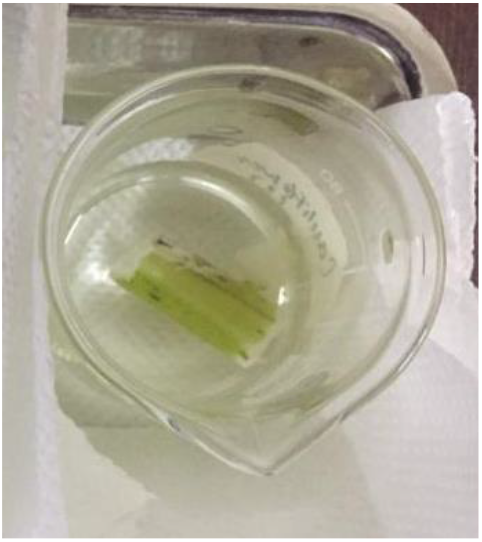
Small cauliflower piece is entirely cleared and large piece is turning white

When both pieces became completely white (took approximately 24 hours for the large piece), we proceeded to the next step of the protocol which involved soaking pieces in tap water for 5 minutes at room temperature to remove any excess bleach. Similar to Protocol A, the cauliflower pieces were then preserved.

### Protocol C

Protocol C is a detergent-based protocol [9]. This protocol used a series of baths to remove nuclear and cellular material, incorporating widely used techniques to decellularize both mammalian and plant tissue [7], [15], [16], [17], [18]. Similar to the first two protocols, two cauliflower pieces of sizes 2.5 cm x 1.1 cm and 1.2 cm x 1 cm were washed with tap water to remove debris and subsequently submerged in tap water on low shake for 20 mins at room temperature. The cauliflower pieces were then placed in a 10% SDS solution for 10 days on low shake at room temperature. In this case, the duration was doubled as cauliflower stalk was thicker than the leaf sample used in the original experiment. It was observed that after 10 days the solution turned brown and sticky making it difficult to remove the cauliflower pieces as shown in Fig. 4.

**Fig 4:**
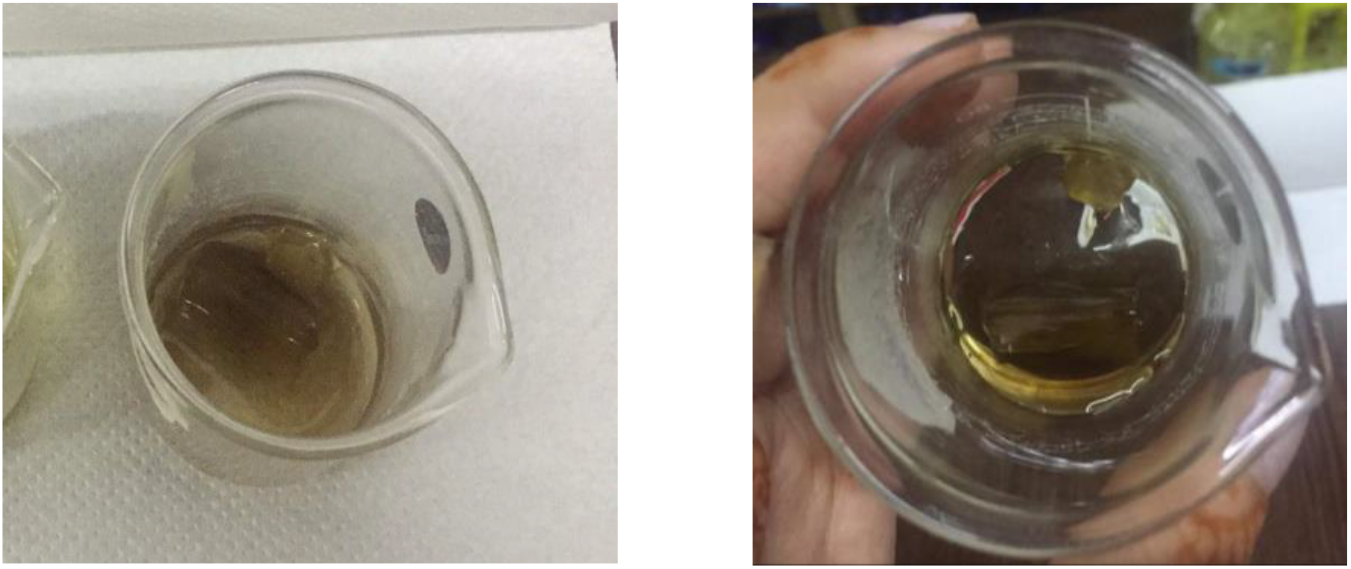
SDS solution turned brown and sticky after 10 days, cauliflower pieces stuck within

Cauliflower pieces were taken out of the viscous 10% SDS solution and washed. The specimen was incubated in tap water for an additional 10 minutes on low shake to remove any residual SDS solution. They were then submerged in a solution of 1% Triton X-100 (the protocol required a 1% non-ionic surfactant) and 5% bleach solution (due to non-availability of 10% bleach as specified in the protocol). Triton X-100 solution was prepared by adding 1mL Triton X-100 to 99 mL cooled boiled water. Cauliflower pieces were then left in the solution on low shake at room temperature until visibly cleared. After becoming fully white, they were transferred to cool boiled water and submerged in it on low shake at room temperature for 10-15 min to rinse off any remaining Triton X-100 and Bleach. Fig. 5 shows the preserved state of the specimen.

**Fig 5:**
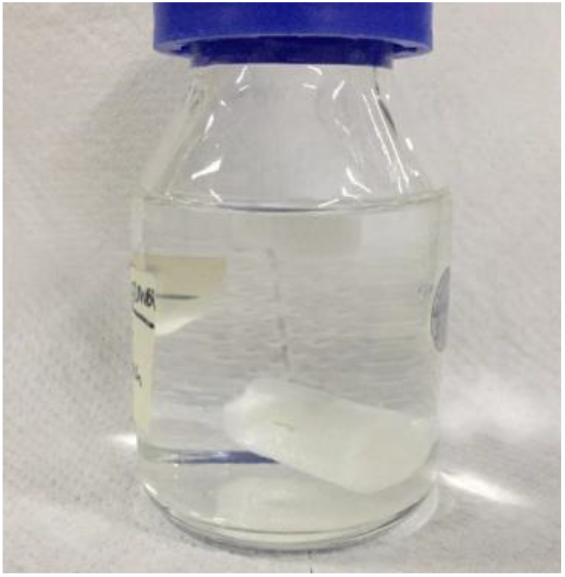
Cleared cauliflower pieces preserved in isopropanol

### Protocol D

We proceeded to follow another protocol that closely resembles Protocol C [10]. The only additional step was in the beginning using n-hexane and 1 x PBS to remove the cuticle. We prepared 1x PBS by dissolving one tablet in 100mL water and filter sterilizing it. The materials and methods [10] do not mention the ratio of volumes of n-hexane and 1x PBS or the duration of soaking. In this protocol, cauliflower pieces measuring 4 cm x 1 cm and 1.9 cm x 0.9 cm were soaked in distilled water for 10 mins to remove debris. We then transferred them to a solution containing 40mL n-hexane and 5mL 1xPBS. We let it soak for 24 hours, only to find no apparent change to the cuticle and the solution to be volatile (see Fig. 6). We decided not to perform the rest of the protocol due to ambiguity in the ratios of volume and duration of submersion.

**Fig 6:**
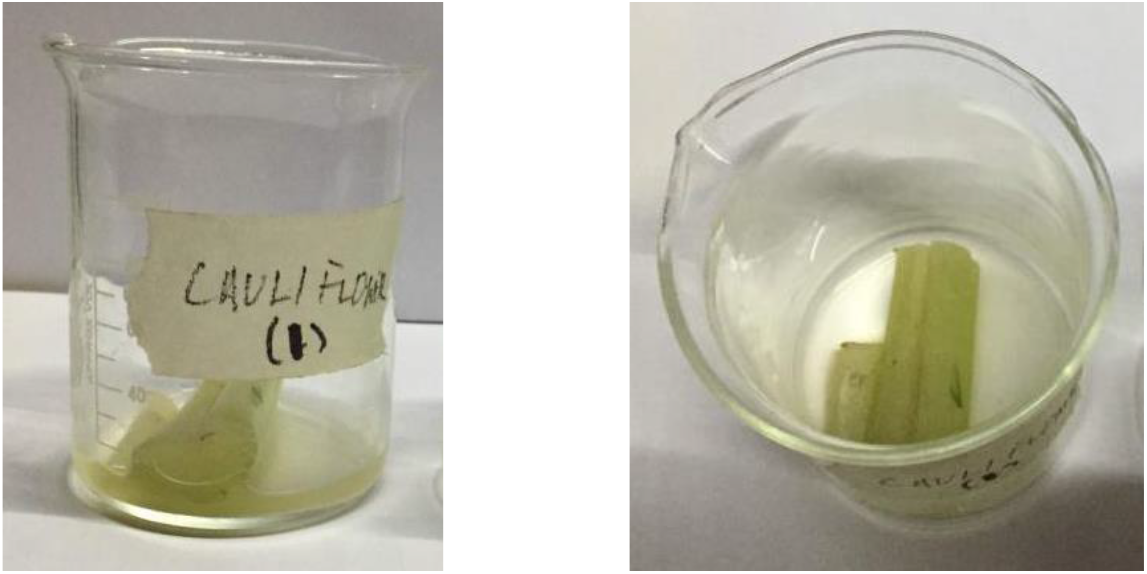
Cauliflower stalks after leaving them in n-hexane and PBS solution for 24 hours

## IV. PROPOSED HYBRID PROTOCOL

Based on the lessons learnt from the experiments using the four protocols, we propose a hybrid protocol and term it as Protocol E. The hybrid protocol was developed on the basis of other protocols through trial and error is discussed here in detail.

### Protocol E

The cauliflower pieces (measuring 2.2cm x 1cm and 0.9 cm x 0.5 cm) were washed and incubated in cool boiled water for 15 minutes on low shake at room temperature to remove any debris. Meanwhile, a solution of 3% Sodium Bicarbonate and 5% Bleach (NaClO) was prepared by using equal volumes of both. The solution was then warmed to 70°C in a water bath. They were then submerged in the prepared solution on low shake at room temperature until visibly cleared. Once they become fully white, they were incubated in cool boiled water for 10 minutes on low shake to remove any residual solution. We then proceeded to incubate the pieces in a 10% SDS solution for approximately 18 hours on low shake after which they were submerged in cool boiled water for another 10 minutes to remove any excess SDS solution before being preserved in isopropanol for microscopy.

## V. HISTOLOGICAL ANALYSIS

We performed microscopy on cauliflower pieces that have been treated through various decellularization protocols through an inverting microscope of 10x, 20x and 40x magnification. We used Safranin dye to stain the nuclei dark red. Absence of dark nuclear stains confirmed the absence of nuclear material and hence the success of the decellularization process.

For microscopy, preserved cauliflower specimens were cut into thin pieces using a dissecting microscope and is flooded with distilled water. These small pieces were then stained with Safranin dye for 3-5 minutes before being washed with distilled water for an additional minute. Cauliflower pieces were then mounted on a slide and observed with an inverted microscope.

As a control, we used an untreated cauliflower specimen for comparison purposes. A thin sample was washed, dyed and observed under an inverted microscope at 10x and 40x magnification.

Fig. 7 shows two magnified views of the specimen. In Fig.7 (b), the edge of the specimen showed dense nuclear material in the central and lower regions of the slide. We then took a thicker specimen from the same sample of untreated cauliflower and repeat the process. Fig. 8 shows a 40x magnification of the thick control specimen from either ends. Dense regions were visible, indicating the presence of nuclear material. All other samples that we further observed were compared to the control specimen to draw a visual comparison and assess the extent of decellularization. The same dying procedure was applied.

**Fig 7:**
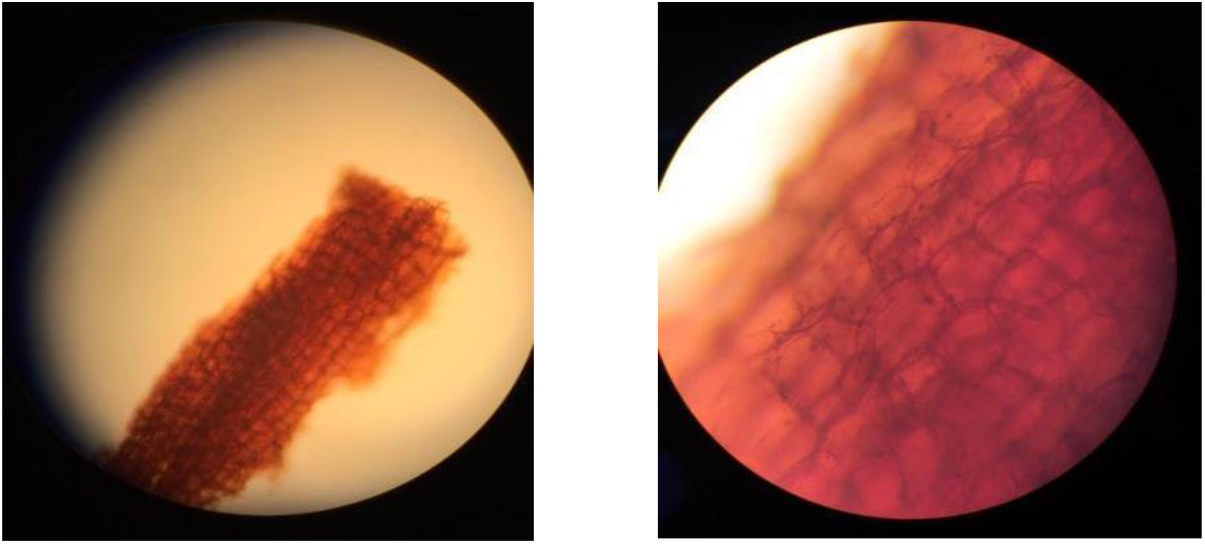
(a) Control specimen under inverted microscope 10x (b) 40x

**Fig 8:**
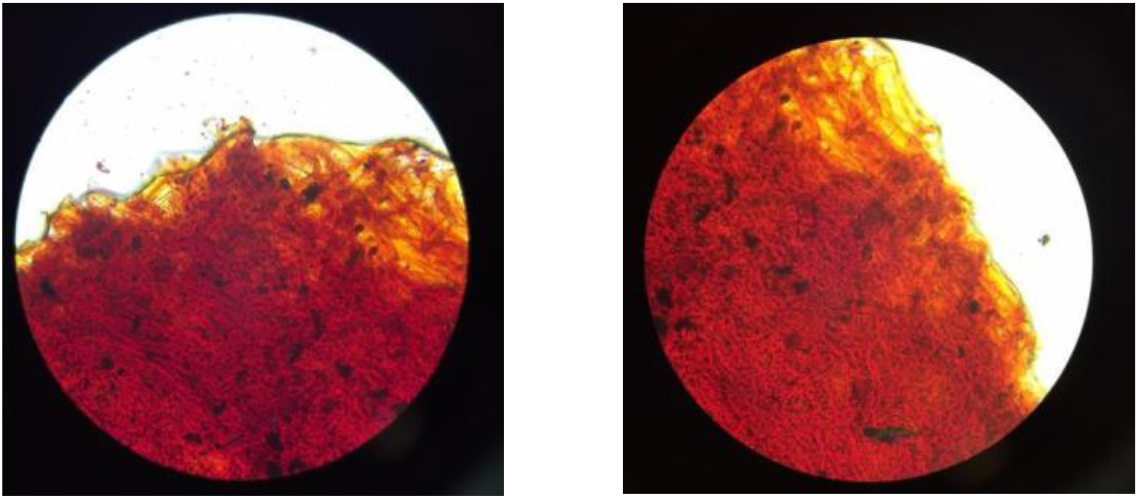
Dense regions in control specimens observed at 40x.

A specimen from Protocol A was observed at 10x, 20x and 40x (see Fig. 9). The central region was magnified at 40x (Fig. 9 c), the cells appeared to be empty and no dense nuclear material was observed.

**Fig 9:**
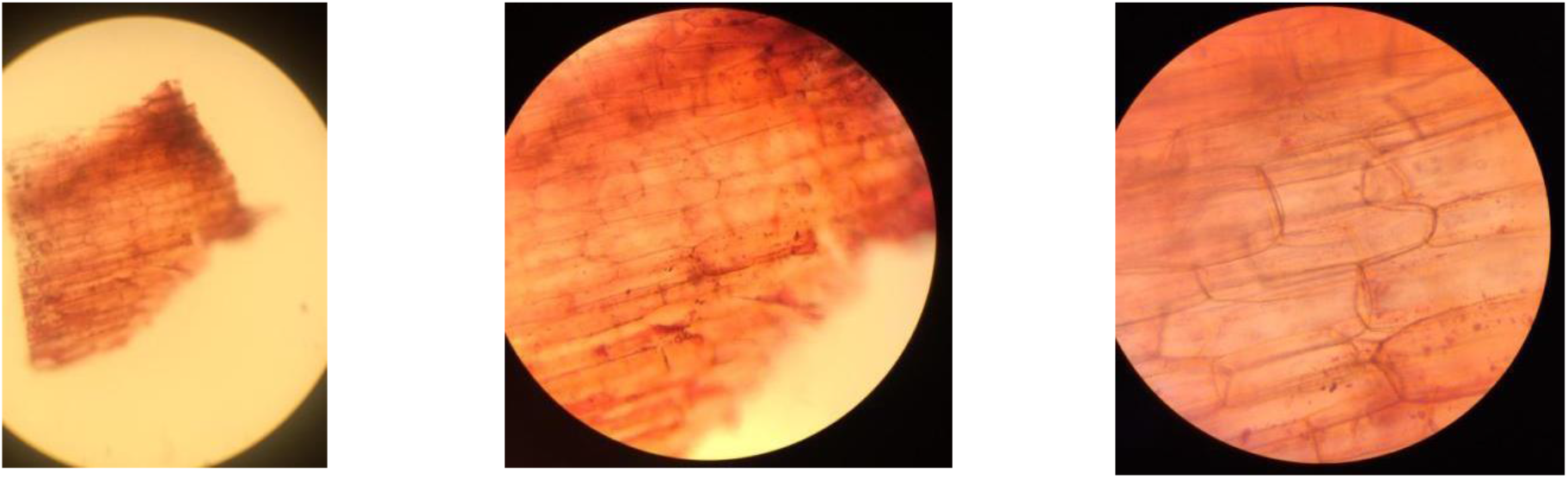
Cauliflower pieces from protocol A under inverted microscope 10x (left), 20x (middle) 40x (right)

Specimen from Protocol B was observed at 40x magnification as shown in Fig. 10. Denser regions were seen and the tissue appeared to be darker. Nuclear material was observed in many cells, either due to partial or incomplete decellularization altogether.

**Fig 10:**
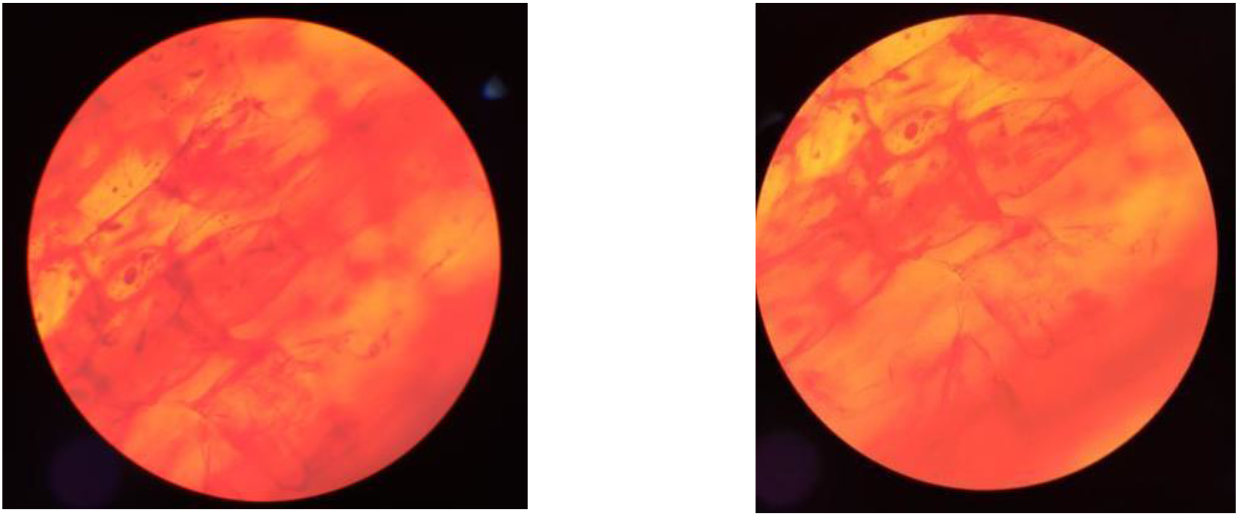
Cauliflower pieces from protocol B under inverted microscope, 40x

Fig. 11 shows the extent of decellularization of Protocol C. The lower end of the specimen was observed under 40x magnification. This enabled us to observe dense tissue and even denser regions of nuclear material within cells.

**Fig 11:**
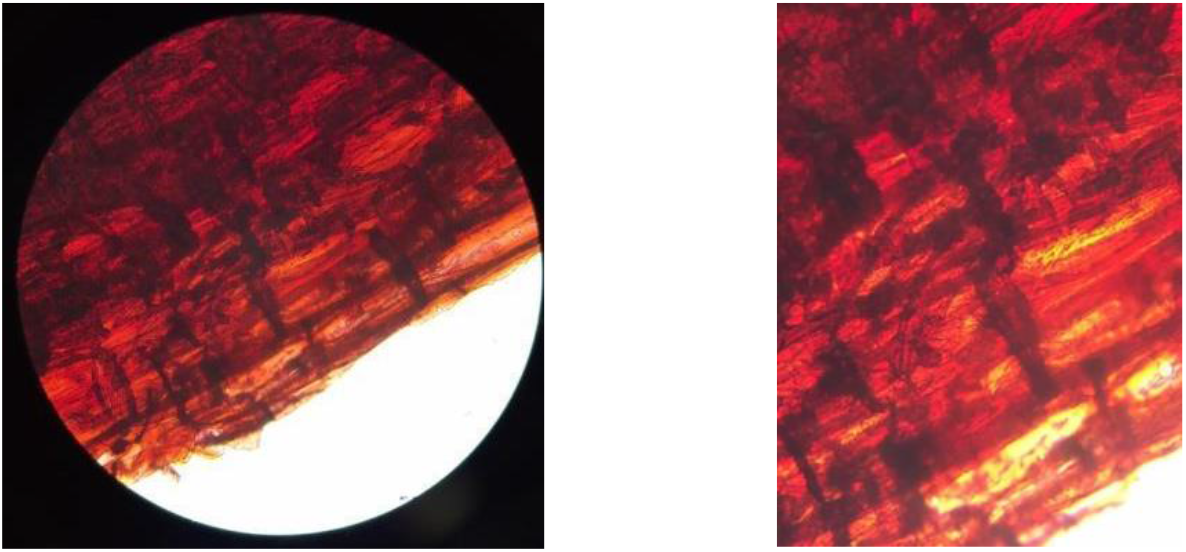
Cauliflower pieces from protocol C observed at 40x

Finally, we analyzed specimens treated with Protocol E. The central and peripheral regions of the specimen were observed under 40x magnification in Fig. 12. It showed cells that are visibly empty, no dense regions can be seen. Out of all slides observed, these appeared to be the one showing maximum decellularization.

**Fig 12:**
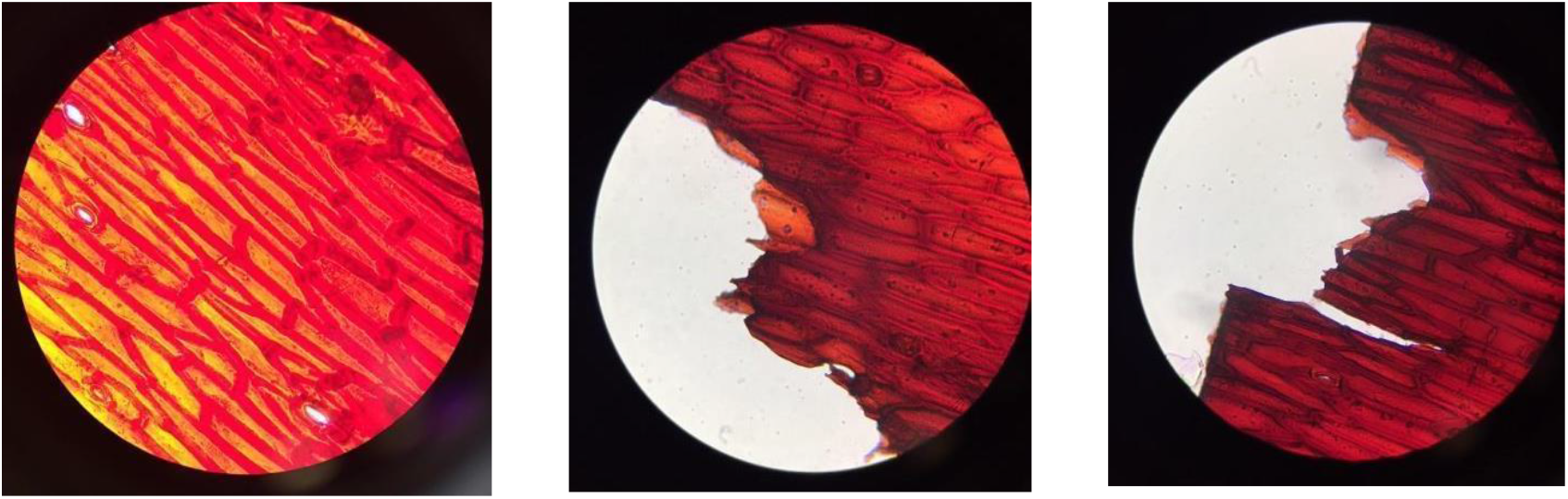
Cauliflower pieces from Protocol E at different sites using inverted microscope (40x)

## VI. RESULTS

The microscopy results of each protocol were compared to the control specimen. Histological analysis of our specimens showed that the cauliflower pieces obtained from Protocols B and C yielded results that resemble the control specimen. In Fig. 10, nuclear material was observable in the cells, suggesting that decellularization did not take place. The same could be inferred for Protocol C as Fig. 11 shows dense tissues with intact nuclear material. At this stage, we could not ascertain any definitive reason as to why these two protocols failed. We anticipated that for Protocol C, it was possible that due to slight changes in duration of cauliflower submersion in the 10% SDS solution and use of 5% bleach instead of 10% may have contributed to this outcome.

Protocols A and E yielded the best results. As observed in Fig. 9, lack of densely stained nuclear material was a strong indicator of completion of decellularization. The tissues were also significantly lighter than others. Moreover, cells observed in the specimen from Protocol E were hollow.

### High-Resolution Images

The next phase in this research was the development of computer-aided design (CAD) model from the images of scaffolds. The real challenge was posed by the internal surface of the scaffold that required high-quality resolution of the images. A digital camera of 24.2 mega pixels was used to take multiple snapshots at various angles that could assist in the development of (CAD) model of the specimen. Cauliflower pieces from the protocols that yielded the best results (Protocols A and E) were prepared to be designed and sent for 3D printing. Fig. 13 shows different images of the specimen.

**Fig 13:**
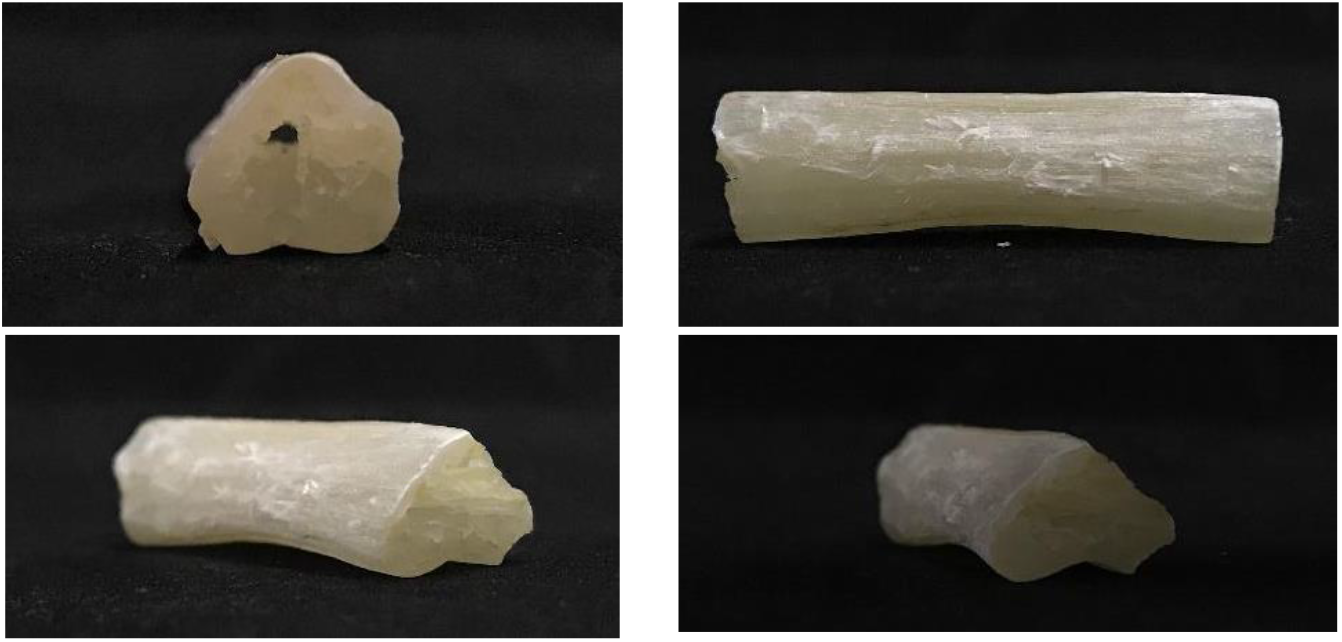
Different images of the scaffold from several angles

### Scaffold CAD Model

The scaffold design was developed from the images using CAD software SolidWorks [20]. In order to trace the surface contours of the scaffolds, B-splines were used. B-splines are more suitable results for complex curves and minimize errors at every cross-section of the scaffold slice. We minimized error to the best of our ability to obtain the CAD model as close to the scaffold as possible. The mean-square error (*MSE*) is defined as follows:

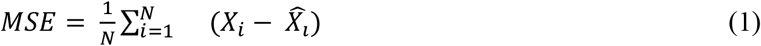

where *X*_*i*_ is the actual coordinate value on the surface from the image and 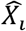is constructed using B-spline. The CAD model allowed us to view different cross-sectional views of the scaffold which was not possible without intrusive dissection of the scaffold. This feature of the model can be exploited for detailed analysis. Fig. 19 showed different views of the scaffold including the cut-away model along the axial axis showing the inner curvature along the length. The outer profile provided a good comparison to the original images shown in Fig. 18.

**Fig 19:**
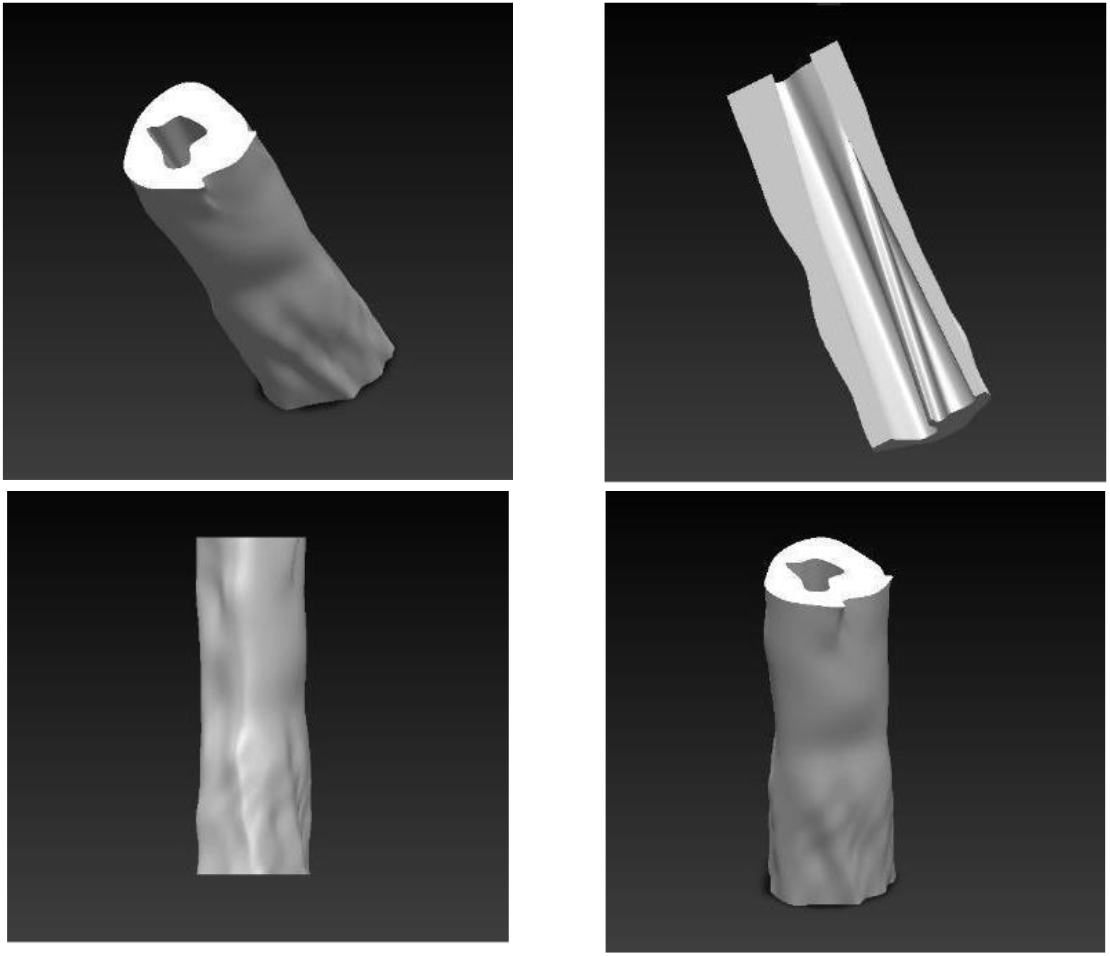
3-D CAD model of the scaffold with different views

### 3-D Printing

The CAD files developed from the images of scaffold were saved as .STL files with fine resolution that resulted in an STL file with ideal accuracy for most 3D printing applications. STL files, (Standard Triangle Language / Standard Tessellation Language) described only the surface geometry of a three-dimensional object without any representation of color, texture or other common CAD model attributes and is a standard format for 3D printing. Using high resolution, we printed the 3-D structure of the scaffold using a XYZprinting da Vinci Series Full Color 3-D printer with PLA filament. The 3-D model mirrored all grooves and imperfections of the scaffold. We printed two sizes from the same model, highlighting the advantage of scalability.

**Fig 20:**
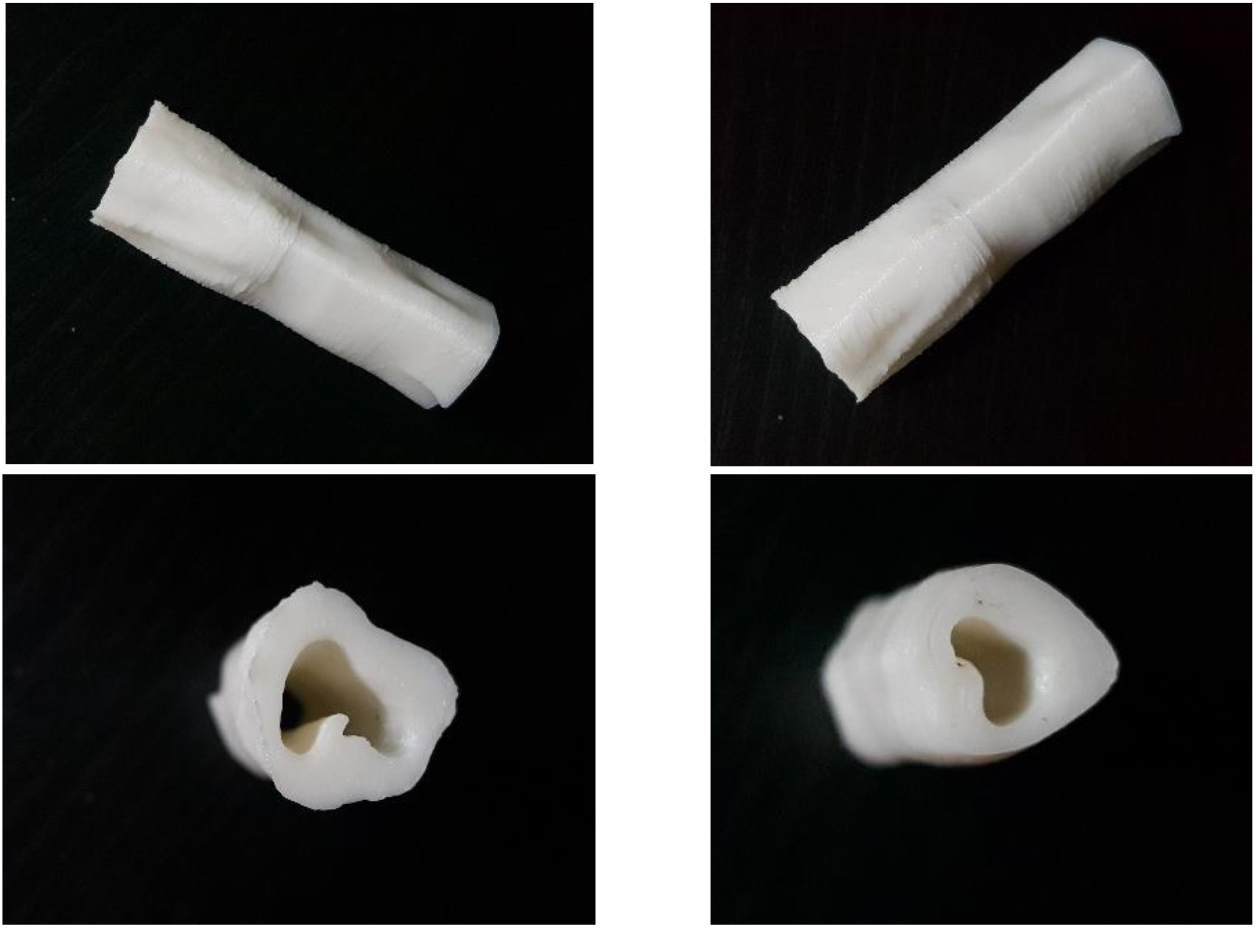
Different views of 3-D printed scaffold using PLA filament

## VII. DISCUSSION AND CONCLUSION

Based on our experience with different protocols for decellularization, we propose a hybrid protocol. A solution of 3% Sodium Bicarbonate and 5% Commercial Bleach followed by 10% SDS Solution yielded the best results as presented in Section V. There is clearly a substantial room for improvement of the decellularization step, more specifically for stems. For example, the width and density of the choice of specimen is one. Most papers that have been published focus on less dense tissues such as leaves, which albeit have a flat architecture unlike blood vessels, but might be easier to decellularize. This could be one of the reasons why some protocols failed. Most papers lack complete details about their protocols which we found to be a hindrance in reproducing them. For example specifications including duration of some steps the ratio of volumes for preparing certain solution are not always mentioned. Finally, the ideal experiment for verifying our deceullarization results could have been the use of DNA quantification using DNA extraction methods followed by gel electrophoresis or flourometry, however due to time constraints we were only able to perform histological analysis.

In future attempts, we hope to use ‘bioinks’ and use new 3-D Bio printing technology instead of the regular plastic 3-D printing. This will allow us to construct a scaffold of our choice using appropriate biological materials to experiment with the recellularization process. This will be tailored to the need and specificity of the organ/vessel needed to be reconstructed.

Through bioprinting of an existing image, direct cellularization of a printed scaffold would become possible without having to carry out the decellularization protocol each time, expediting the iteration process. This would also remove the need to carry out any analysis each time to verify the quality of decellularization save more time and cost. Finally, the highest impact of such a “decellularize, image, print” cycle is the ability to ‘capture’ scaffold structures from different plant sources and digitally share them with other, innovators and DIY biologists who can use a scaffold for any experiment right away. These could digital models of scaffolds can be easily scaled to the desired size, readily shared between labs and open sourced in a public repository. By increasing the pace of innovation in the tissue engineering space, this has the potential to develop standard templates for use as scaffolds for particular organs, tissues or other parts and potentially revolutionize tissue engineering.

## ACKNOWLEDGEMENTS

The first author would like to thank Pakistan Innovation Foundation for the STEMx Internship Program that provided the opportunity to work with Dr Faisal Khan. The authors are indebted to Mr. Usman Akhtar for his assistance in 3D printing of the specimen model.

